# Assembling highly repetitive *Xanthomonas* TALomes using Oxford Nanopore sequencing

**DOI:** 10.1101/2022.08.17.504259

**Authors:** Annett Erkes, René Grove, Milena Žarković, Sebastian Krautwurst, Ralf Koebnik, Richard D. Morgan, Geoffrey G. Wilson, Martin Hölzer, Manja Marz, Jens Boch, Jan Grau

**Affiliations:** Institute of Computer Science, Martin Luther University Halle-Wittenberg, Halle, Germany; Department of Plant Biotechnology, Leibniz Universität Hannover, Hannover, Germany; Bioinformatics/High-Throughput Analysis, Friedrich Schiller University Jena, Jena, Germany; PHIM Plant Health Institute, Univ Montpellier, IRD, CIRAD, INRAE, Institut Agro, Montpellier, France; New England Biolabs Inc., Ipswich, MA, USA; Methodology and Research Infrastructure, MF1 Bioinformatics, Robert Koch Institute, Berlin, Germany

## Abstract

Most plant-pathogenic *Xanthomonas* bacteria harbor transcription activator-like effector (TALE) genes, which function as transcriptional activators of host plant genes and support infection. The entire repertoire of up to 29 TALE genes of a *Xanthomonas* strain is also referred to as TALome. The DNA-binding domain of TALEs is comprised of highly conserved repeats and TALE genes often occur in gene clusters, which precludes the assembly of TALE-carrying *Xanthomonas* genomes based on standard sequencing approaches. Here, we report the successful assembly of the 5 Mbp genomes of five *Xanthomonas* strains from Oxford Nanopore Technologies (ONT) sequencing data. For one of these strains, *Xanthomonas oryzae* pv. *oryzae* (*Xoo*) PXO35, we illustrate why Illumina short reads and longer PacBio reads are insufficient to fully resolve the genome. While ONT reads are perfectly suited to yield highly contiguous genomes, they suffer from a specific error profile within homopolymers. To still yield complete and correct TALomes from ONT assemblies, we present a computational correction pipeline specifically tailored to TALE genes, which yields at least comparable accuracy as Illumina-based polishing. We further systematically assess the ONT-based pipeline for its multiplexing capacity and find that, combined with computational correction, the complete TALome of *Xoo* PXO35 could have been reconstructed from less than 20,000 ONT reads. Our results indicate that multiplexed ONT sequencing combined with a computational correction of TALE genes constitutes a highly capable tool for characterizing the TALomes of huge collections of *Xanthomonas* strains in the future.

## INTRODUCTION

*Xanthomonas oryzae* plant-pathogenic bacteria cause highly devastating diseases of the major food crop rice. *X. oryzae* pv. *oryzae* (*Xoo*) and *X. oryzae* pv. *oryzicola* (*Xoc*) are two related groups of these pathogens which differ by their mode of infection. While *Xoo* spreads via the vascular systems, *Xoc* predominantly infects the parenchyma tissue of the rice leaves (1). The virulence of both pathovars is dependent on a type-III secretion system which injects a collection of different effector proteins directly into plant cells to manipulate and reprogram these to support the bacterial infection (2).

Transcription activator-like effectors (TALEs) constitute the largest effector family in *X. oryzae* with up to 19 (*Xoo*) and 29 (*Xoc*) different TALEs within one strain (3–5). TALEs function inside the plant nucleus as transcription factors to induce transcription of specific target genes (6, 7). The outstanding feature of TALEs is their modular DNA-binding domain, consisting of a variable number of 34 amino acid-repeats with each repeat facilitating binding to one DNA base in a sequential fashion (8, 9). The amino acid 13 in each repeat directly contacts the base in the leading strand of the double-stranded DNA and the combinations of amino acids 12 and 13 in each repeat form repeat-variable diresidues (RVD), which determine the binding preference of an RVD and, consequently, the DNA-binding specificity of each TALE protein (Fig. 1A) (8–11). TALEs interact with subunits of the transcription initiation factor IIA (12) and initiate transcription via a C-terminal acidic activation domain in a somewhat defined distance downstream of the TALE binding site (13, 14). Accordingly, bioinformatic prediction of TALE target sites in plant promoters in combination with gene expression analysis of host plant genes after *Xanthomonas* infection is the standard procedure to identify potential virulence targets of TALEs (4, 7, 15, 16).

**Fig. 1.**
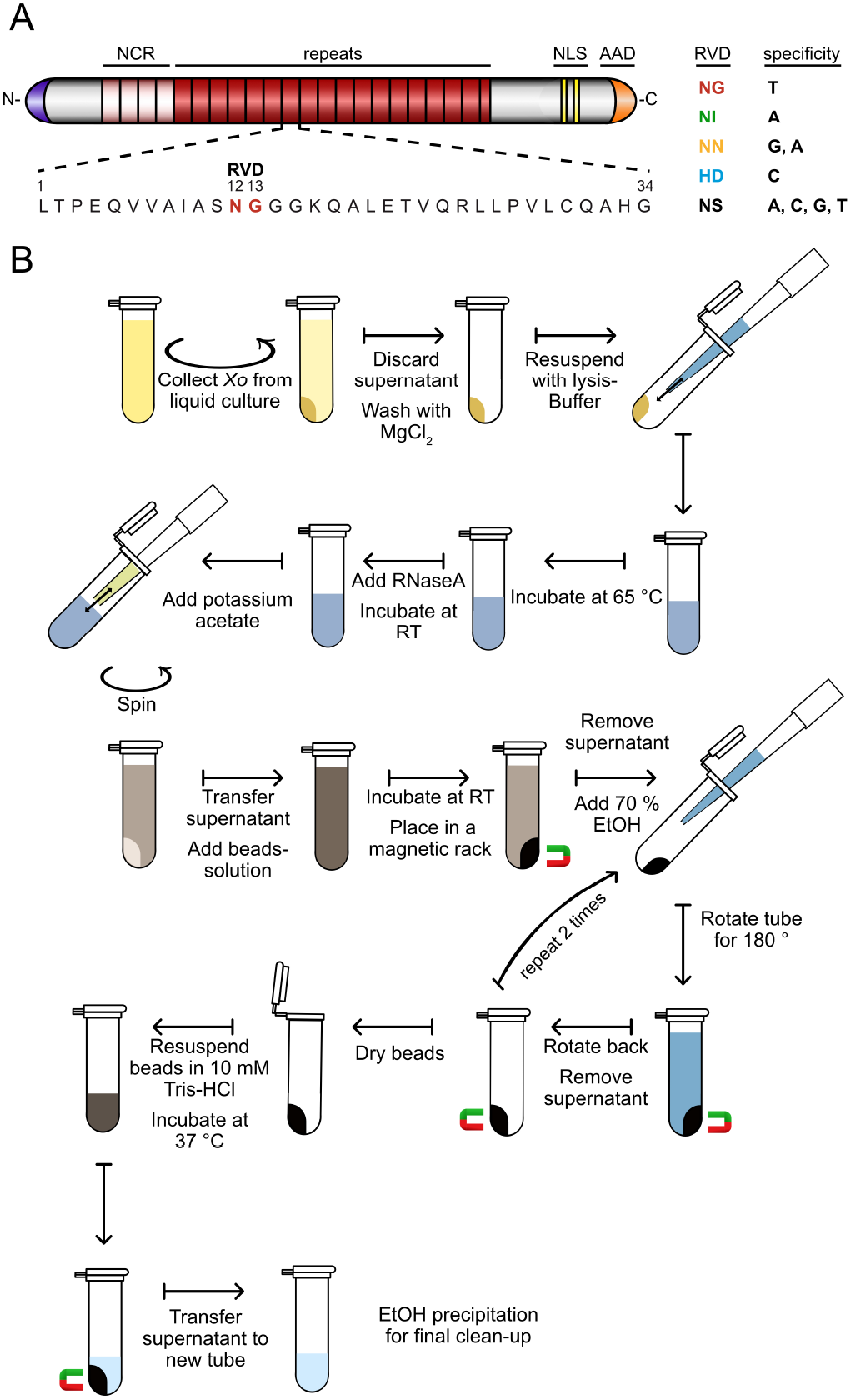
(A) Schematic representation of the structure of a TAL effector with four non-canonical repeats (NCR, light red), tandem repeats (red), nuclear localisation signals (NLS) and acidic activation domain (AAD). The 34 amino acids of each repeat are highly conserved, excluding the amino acids at position 12 and 13, which are referred to as repeat variable diresidue (RVD). (B) A magnetic bead-based method to isolate high molecular weight genomic DNA for ONT sequencing.

Different strains of *Xoo, Xoc* and other Xanthomonads carry individual repertoires of TALEs which evolve by base substitution in RVD codons, recombination between TALEs and deletion of repeats within a TALE (17–20). TALEs and other effector genes in *Xanthomonas* species are often flanked by 62 bp inverted repeats (21) suggesting that they can be mobilized by transposition (22). The structures of TALE gene clusters in *Xoc* additionally suggest an integron-like mechanism for the reassortment of TALEs (18). Functional TALEs contain between 7.5 and 33.5 repeats (6) and this repetitiveness precludes the correct assembly of TALE genes using short read-based next generation sequencing techniques. Accordingly, assemblies of *Xanthomonas* genome sequences following Illumina sequencing typically exclude the TALEs (23). The first complete *Xoo* (24) and *Xoc* (25) genome sequences including TALEs were produced using Sanger sequencing on subcloned genomic fragments, but this approach is now outdated as it is time-consuming and not feasible for high throughput. Shortcuts to determine the set of TALEs (e.g., the TALome) within a strain used either direct subcloning of conserved restriction fragments (26, 27), Gibson assembly of these fragments (28) or PCR to amplify repeat regions of TALEs and subsequently subcloning and sequencing those (29). However, these approaches omit the information how TALEs are organized in clusters and risk to overlook individual TALEs.

The current gold standard for sequencing *Xanthomonas* genomes including *TALE* genes is PacBio sequencing, which can produce high-quality, circular, single contig genome assemblies (4, 17, 19, 30–32). Nevertheless, this technique requires complex protocols and expensive platforms compared to standard short-read technologies. Furthermore, some *Xoc* TALE gene clusters contain ten consecutive TALE genes, reaching a length of 40.7 kbp (*Xoc* CFBP2286) (3) which corresponds to only the top five percent of the typical PacBio read length (33). Also, genomes of *Xanthomonas* (as well as those of other plant pathogen bacteria) contain high numbers (up to ten percent) of mobile elements including insertion sequences (IS), transposons and phage-related genes which trigger frequent genomic rearrangements (22, 34, 35). Duplication of TALE clusters as well as rearrangements of TALEs between clusters by inversions can lead to repetitive genomic structures that are not easily resolved.

In general, decoding the correct arrangement of repetitive elements in genomes has been a long-standing challenge for sequencing techniques. A notorious example for this are the centromeric and telomeric regions of eukaryotic genomes (36). The ultra long reads (*>* 100 kbp) of single DNA molecules obtained from Oxford Nanopore Technologies (ONT) sequencing now finally provide a solid approach to this problem, although the high error rate of 5–15 percent is currently still a challenge (37). Combining these long reads with circular PacBio HiFi reads or Illumina reads can significantly increase the read accuracy while solving the genome structure at the same time (38–41).

Here, we report the establishment of an ONT sequencing approach for *Xoo* and *Xoc*, and a specific computational read assembly pipeline including an *in silico* correction method to increase the accuracy of TALE gene assemblies. With this, we were able to solve the arrangement of a particularly complex genomic TALE cluster in *Xoo* strain PXO35, which was not possible for us based on PacBio sequencing data. Contiguity and genome completeness were evaluated for assemblies based on Illumina, PacBio and ONT sequencing data, and based on hybrid approaches, respectively. The possibility to multiplex this approach will enable sequencing and characterization of TALomes of multiple *Xanthomonas* strains in parallel on a single ONT flow cell, and will allow the identification of evolutionary events within *Xoo* and *Xoc* populations.

## METHODS

### Bacterial growths

*Xanthomonas oryzae pv. oryzae* (*Xoo*) strains PXO35, FXO38 and Huang604 and *Xanthomonas oryzae pv. oryzicola* (*Xoc*) strains BAI35 and MAI23 were cultivated in PSA medium at 28 °C.

### DNA extraction and library preparation

#### Illumina

Bacteria were washed in 1M NaCl and genomic DNA was isolated using the Wizard® genomic DNA purification kit (Promega, Charbonnières, France).

#### PacBio

Genomic DNA from *Xoo* PXO35 was isolated using the DNeasy Blood & Tissue kit with pretreatment for Gram-negative bacteria according to the protocol from QIAGEN.

#### Production of Magnetic Nano Particles

Magnetic Nano Particles (MNPs) were made and silica-coated according to Oberacker et al. (2019)(42).

#### gDNA Extraction from *Xanthomonas oryzae* using Magnetic Nano Particles

In the following, we describe the protocol for gDNA extraction using magnetic nano particles (MNPs), which is also illustrated in Figure 1B. Additional details about the preparation of the beads solution is provided as Supplementary Methods.

First, 4 ml *Xoo*/*Xoc* overnight culture were collected by centrifugation (5 min, 10,000 g) in a 2 ml reaction tube. The pellet was washed by resuspension in 1 ml of 10 mM MgCl2 and pelleted again by centrifugation. The pellet was then resuspended in 900 μl pre-heated lysis buffer (0.5 M NaCl; 100 mM Tris-HCl, pH 8.0, 50 mM EDTA; 1.5 % SDS; 65 °C) by pipetting up and down. The mixture was incubated at 65 °C for 20 min with shaking of 500 rpm. After 10 min the tube was inverted for 10 times. 4 μl RNase A (10 mg/ml; DNase free) was added and incubated for 10 min at room temperature. After RNA digestion, 300 μl potassium acetate (5 M) was added and mixed by pipetting up and down with a 1,000 μl pipet tip, which was cut 0.5 cm from the tip end. For pelleting the insoluble compounds, the mixture was centrifuged for 10 min at 20,000 g. 800 μl of the supernatant was transferred to a new 2.0 ml tube, mixed by inverting with 800 μl of well homogenized beads solution and incubated for 1 h with gentle shaking. Tubes were then placed on a magnetic rack until all beads were stuck to the tube wall. The supernatant was discarded and the beads were washed in 70 % EtOH for 3 times, by adding the EtOH to the tube with the beads, and rotating the tube for 180° while it stays in a magnetic rack. After a second rotation for 180 ° and a complete binding of the beads to the tube wall, the EtOH was removed by pipetting. After washing, remaining EtOH was removed by incubation for 5 min at 37 °C with an open lid. To elute the DNA from the beads, beads were resuspended in 100 μl 10 mM Tris-HCl, pH 8.5 and incubated at 37 °C with shaking of 500 rpm for 60 min. Using a magnetic rack, the supernatant containing DNA was seperated from the MNPs into a new 1.5 ml tube. By adding 1/10 Volume of 5 M potassium acetate and 2 volumes of EtOH, a precipitation of the DNA was performed. After an incubation of 30 min at room temperature, the precipitated DNA was centrifuged for 10 min at 20,000 g. The supernatant was removed and the pellet was washed with 70 % EtOH for two times. After washing, the remaining EtOH was removed by incubation at 37 °C for 5 min with an open lid. The DNA was resuspended in 50 μl 10 mM Tris-HCl (pH 8.5) by shaking (500 rpm) for 1 hour at 37 °C.

### Sequencing

#### Illumina sequencing of *Xoo* PXO35

Strain PXO35 was sequenced using the Illumina HiSeq 2500 platform (Fasteris SA, Switzerland). The shotgun sequencing yielded 2,978,197 100-bp paired-end reads (596 Mb), with insert sizes ranging from of 250 bp to 1.5 kb.

#### PacBio sequencing of *Xoo* PXO35

Libraries for *Xoo* PXO35 were sequenced in two Pacific Biosciences SMRT cells as described previously (4), yielding a total of 143,946 continuous long reads (CLRs) with an N50 read length of 21,760 bp.

#### Oxford Nanopore

Library was made using Oxford Nanopore Technologies protocol: Native barcoding genomic DNA with EXP-NBD104 and SQK-LSK109 kit. Manufacturer’s instructions were followed with slight changes. 700 ng of each end-prepped sample was barcoded with barcode (1-FXO38, 2-PXO35, 3-MAI23, 4-Huang604, 5-BAI35). Incubation time for end-prep reaction was prolonged to total of 30 minutes, and elution was 5 minutes on 37 degree. 150 ng of each barcoded sample was pooled together (total 750ng) for adapter ligation reaction, and here also incubation was prolonged to 15 minutes, and elution to 20 minutes on 37 degree to improve the recovery of the long fragments. Total of 476 ng library was loaded without loading beads to R9.4 flow cell. Sequencing was performed for all samples (5 isolates) on a MinION Mk1B device and basecalled with Guppy v4.0.11 and the dna_r9.4.1_450bps_modbases_damdcm-cpg_hac.cfg model. Parameters were specified for an increased chunk size of 10000 (--chunk_size) and to enable barcode demultiplexing according to the used kit (--barcode_kits).

General statistics of all sequenced libraries are given in Table 1. Although the N50 read lengths of the PacBio and ONT libraries are similar, the ONT library contains a large fraction of very long reads that are less present in the PacBio data. On the contrary, about half of the ONT reads but only approx. 5% of the PacBio reads are shorter than 1 kbp (cf. Supplementary Table A, Supplementary Figure S1).

**Table 1.**
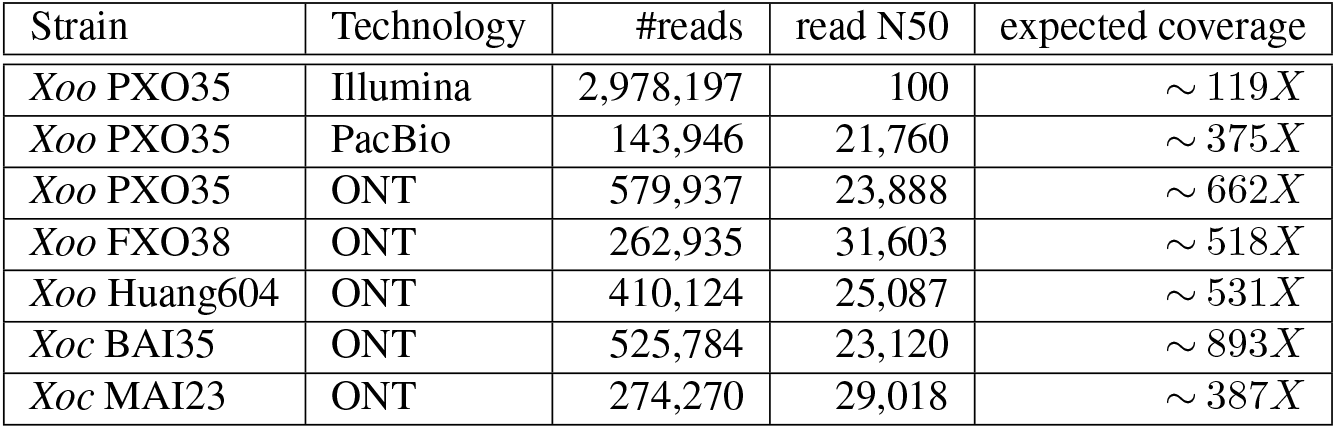
Sequencing statistics of libraries

**Table 2.**
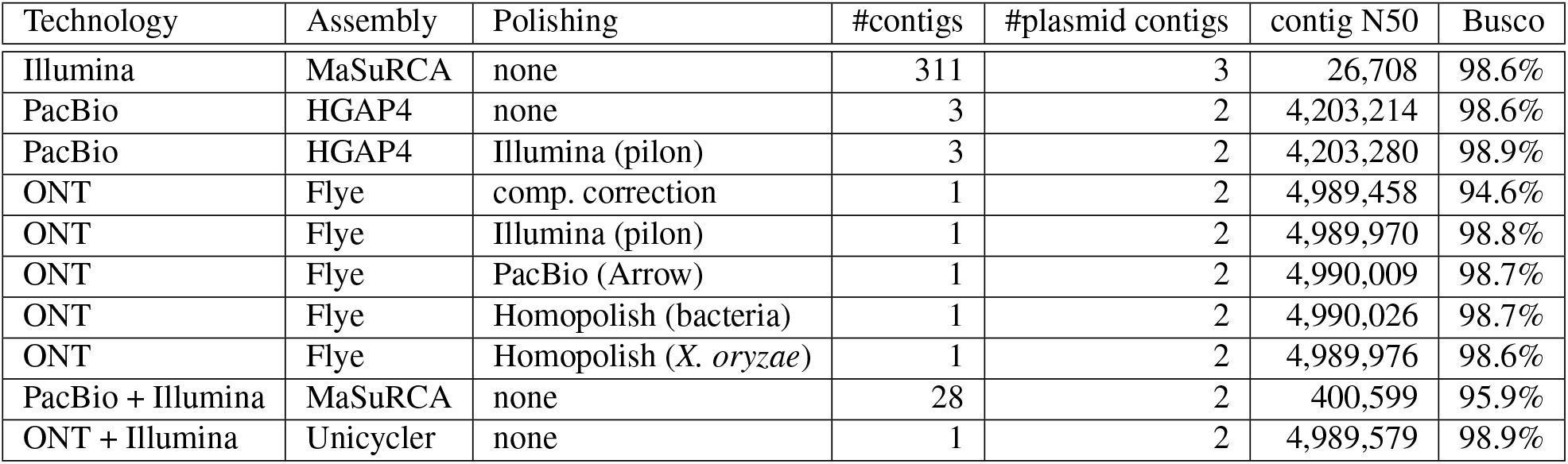
General statistics of genome assemblies of PXO35 using different combinations of input reads, assembly tools and polishing methods including computational correction of TALE genes (comp. correction), listing the number of contigs (#contigs), additional contigs that are likely related to plasmid sequences (#plasmid contigs), the contig N50 value, and Busco completeness per assembly.

| Technology | Assembly | Polishing | #contigs | #plasmid contigs | contig N50 | Busco |
| --- | --- | --- | --- | --- | --- | --- |
| Illumina | MaSuRCA | none | 311 | 3 | 26,708 | 98.6% |
| PacBio | HGAP4 | none | 3 | 2 | 4,203,214 | 98.6% |
| PacBio | HGAP4 | Illumina (pilon) | 3 | 2 | 4,203,280 | 98.9% |
| ONT | Flye | comp. correction | 1 | 2 | 4,989,458 | 94.6% |
| ONT | Flye | Illumina (pilon) | 1 | 2 | 4,989,970 | 98.8% |
| ONT | Flye | PacBio (Arrow) | 1 | 2 | 4,990,009 | 98.7% |
| ONT | Flye | Homopolish (bacteria) | 1 | 2 | 4,990,026 | 98.7% |
| ONT | Flye | Homopolish (*X. oryzae*) | 1 | 2 | 4,989,976 | 98.6% |
| PacBio + Illumina | MaSuRCA | none | 28 | 2 | 400,599 | 95.9% |
| ONT + Illumina | Unicycler | none | 1 | 2 | 4,989,579 | 98.9% |

### Genome assembly

We used matching assembly tools to generate genome assemblies of reads obtained from the different sequencing methods. We assembled ONT reads of *Xoo* PXO35, FXO38 and Huang604 and *Xoc* BAI35 and MAI23 each with Flye (v2.8.3) (43) using parameters “--plasmids –meta --minoverlap 5000”. For the initial assembly, we filtered reads using Filtlong (v0.2.0, https://github.com/rrwick/Filtlong) with parameter “--target_bases 500000000” to obtain a subset of high-quality reads. Mapping of the complete set of ONT reads to the respective assemblies was performed with minimap2 (v2.17-r941) (44). Assemblies were initially polished using these long-read-mappings and one round of Racon (v1.4.13) (45) followed by medaka (v1.2.6, https://github.com/nanoporetech/medaka).

For all strains, we additionally ran Flye without the “--min-overlap” parameter and proceeded with the assembly with the best contiguity. Only in case of MAI23 the latter works best obtaining 1 instead of 2 contigs.

The origin of replication (ORI) was determined based on the proximity to *dnaA, dnaN* and *gyrB* genes and on the G-C content along the genome. The assemblies were rearranged according to their ORI to compare them using the genomic alignement viewer Mauve 2.4.0 (46, 47). In general, the circular chromosomal contig generated by Flye had sequence sniplets or duplications at the circularisation site, which were not covered by mapped reads.

However, this could be solved by further polishing rounds with medaka: For FXO38 we polished two rounds with medaka and for Huang604 three rounds. For all other strains one polishing round with medaka already solved circularization problems.

For *Xoo* PXO35, we generated genomic Illumina and PacBio sequencing data in addition to the ONT data described previously. For the assembly of Illumina reads, we used MaSuRCA gemome assembler version 4.0.3 (48) with default settings. For the assembly of PacBio reads, we used HGAP4 of SMRT LINK (v.7.0.1) (49) with default settings. For the hybrid assembly of Illumina and PacBio reads, we also used MaSuRCA gemome assembler version 4.0.3 (48) with default settings. For the hybrid assembly of Illumina and ONT reads, we used Unicycler version v0.4.8 (50) with default settings.

### Polishing

#### Illumina-based

For the ONT-based assembly (after initial long-read polishing) and the PacBio-based assembly, we investigated the effect of polishing with Illumina reads using Pilon (v1.24) (51). For this purpose, the reads were mapped to the obtained contigs with BWA-MEM (v0.7.17-r1188) (52).

We also polished the pure Illumina assembly generated with MaSuRCA with Pilon and BWA-MEM mapped Illumina reads.

#### PacBio-based

To polish the ONT-based assembly with PacBio reads, we use the Resequencing application from SMRT LINK (v.7.0.1) with default parameters and pbmm2 for the mapping, and choose Arrow as the polishing algorithm.

The PacBio assembly resulting from HGAP4 is polished with PacBio reads in the same manner.

#### Homopolish

Often, Homopolish (53) is used for polishing an ONT-based assembly if additional PacBio or illumina-reads are not available. It polishes the assembly based on homologous sequences. We polish the assembly of *Xoo* PXO35 with Homopolish using the online available homologous sequence data set for bacteria (http://bioinfo.cs.ccu.edu.tw/bioinfo/mash_sketches/bacteria.msh.gz) with parameters “-s bacteria.msh -m R9.4.pkl” and in a second trial with *Xanthomonas oryzae* as NCBI taxonomic name with parameters “-g Xanthomonas_oryzae -m R9.4.pkl”.

### Comparison of assemblies

To suggest a suitable sequencing method for TALE-containing *Xanthomonas* genomes, we compare the PXO35 assemblies obtained with the different sequencing and assembly methods mentioned above based on the number of contigs, contig N50, BUSCO completeness, as well as class and number of containing TALEs. A suitable candidate contains few contigs, a high contig N50, a high BUSCO completeness and many TALEs belonging to known TALE classes. In addition, we compare the assembly candidates based on a genomic alignment. Furthermore, we compare the ONT-sequenced *Xoo* and *Xoc* strains with each other based on the above mentioned features and additionally include two known representatives from *Xoo* and *Xoc* in the comparison of TALomes.

#### Busco genome completeness

To assess the completeness of the assemblies with regard to the genes they contain, we used BUSCO 5.1.3 (54), which uses universal single-copy orthologs to quantify completeness. We restricted the calculation to the orthologs of the “xanthomonadales_odb10” dataset, which contains 1,152 BUSCO markers. We used BUSCO in genome mode with the parameter “-m genome” and report the proportion of complete BUSCOs of each assembly.

#### Alignment using progressiveMauve

To visualise similarities and differences between the different PXO35 assemblies on a genomic level, we created a genomic alignment of all PXO35 assemblies with progressiveMauve 2.4.0 (47). To compare the newly sequenced and assembled strains of the *Xoo* and the *Xoc* pathovar, we generate corresponding genomic alignments using progessiveMauve to investigate similarities and rearrangements between the genomes.

### TALE annotation using AnnoTALE

For identification and analysis of TAL effectors on the assembled genomes we use the application suite AnnoTALE (v1.5) (4). We use the tool ‘TALE Prediction’ and ‘TALE Analysis’ for the detection of TALEs within the sequenced genomes and for determining the RVD composition of TALEs. In addition to the complete TALE DNA sequences, we receive a FastA file with TALE sequences split into the parts N-terminus, individual repeats and C-terminus.

#### Assignment to TALE classes

To assign predicted TALEs to existing TALE classes or to a new class, we use the AnnoTALE tool ‘TALE Class Assignment’ and obtain systematic TALE names due to the nomenclature based on the RVD sequences of the TALEs as introduced with AnnoTALE (4).

#### Comparison of annotated TALEs

For the different assemblies we compare the detected TALEs, based on the number of complete TALEs, iTALEs, pseudo genes and classes. We consider iTALEs separately, as they contain a truncated C-terminus without activation domain and have a distinct role in pathogenicity by interfering with the disease resistance of the plant (55, 56).

#### Comparison to other strains

With the AnnoTALE tool ‘TALE Class Presence’ one can visualize the TALE class composition of the TALomes of different strains. As a reference for the 5 Xanthomonas strains sequenced with ONT, we choose one known representative for each *Xanthomonas oryzae* pathovar: *Xoo* PXO99A and *Xoc* BLS256.

### Computational correction of TALEs in ONT assembly

Using Oxford Nanopore’s sequencing technology to sequence TALE-containing genomes offers the advantage of long reads covering entire TALE clusters, facilitating assembly into a complete genome. However, accuracy in long homopolymer regions is low, even in case of high coverage (57). In case of low coverage, the lower read accuracy of long-read technologies can also cause a problem in non-homopolymeric regions. We present a method for the computational correction of such sequencing errors within TALE sequences from ONT assemblies. To this end, we learned three separate profile hidden Markov models (HMMs) with HMMER (v3.3.2) (58) based on known TALE sequences from AnnoTALE (v1.5). The first HMM represents the N-terminus, the second one individual repeats, and the third one the C-terminus of a TALE. We trained each HMM separately for *Xoo* and *Xoc* TALEs. We then searched for corresponding TALE parts within each ONT assembly using nhmmer (58). We only considered hits with an average posterior probability (acc) above 0.75 and a bit score above 200 in case of N- and C-terminus and above 45 in case of a repeat hit. We searched for insertions and deletions in comparison to the consensus of the HMMs. Only continuous stretches of insertions are considered, which are at most 3 bp (N- and C-terminus) or 2 bp (repeats) long. Within repeats, the di-codon coding for the RVD is excluded from the correction as i) this di-codon is hyper-variable and, hence, the consensus of the HMM is rather meaningless and ii) otherwise RVDs ‘N*’ with the 13th amino acid missing might be corrected erroneously. In case of an insertion, we distinguish between two cases: (i) the insertion occurs within a homopolymer context with at least five identical nucleotides, which results in the insertion of the homopolymer nucleotide, or (ii) otherwise, the corresponding consensus nucleotide of the HMM is inserted. Inserted bases are lower-cased in the corrected TALE assembly to make the correction identifiable in downstream analyses. As an optional step, we aim to improve correction with a custom substitution polishing. First, we map the ONT reads against the TALE-corrected assembly with minimap2 (v2.17-r941) (44) and then adapt the previous correction if another base is 1.5 times as frequent as the base chosen from the HMM consensus. Here, we count occurrence of each base with igvtools count (v2.5.3) (59).

### Testing multiplexing capacity by sub-sampling

We investigate, which coverage is sufficient to decode the TALome of *Xoo* PXO35 and, hence, how many *Xanthomonas* strains could be multiplexed on a single ONT flow cell, by sub-sampling from the ONT reads. For this purpose we randomly sample 75%, 50%, 25%, 12.5%, 6.25% and 3.12% of the original ONT reads with seqtk sample https://github.com/lh3/seqtk. For each of the generated sub-samples, we generate an assembly with Flye, which we further polish with Racon and medaka as described above for the complete set of ONT reads. Subsequently, we apply the computational correction of TALE genes proposed in this manuscript including custom substitution polishing and finally use AnnoTALE for TALE annotation.

### Availability

The scripts for computationally correcting TALE genes within ONT-based assemblies of *Xoo* and *Xoc* genomes are available from github at https://github.com/Jstacs/Jstacs/tree/master/projects/talecorrect. All assemblies of the *Xoo* PXO35 genome considered in Table 3 and the ONT-based assemblies of the remaining strains (cf. Table 5) are available from zenodo (https://doi.org/10.5281/zenodo.6854866, doi: 10.5281/zen-odo.6854866). Raw reads are available from the European Nucleotide Archive (https://www.ebi.ac.uk/ena) under study accession PRJEB54678.

**Table 3.** Comparison of PXO35 assemblies on level of TALEs. Per assembly, we record the total number of TALE genes (#TALEs), the number of iTALEs with a truncated C-terminus (#iTALEs), the number of TALE pseudo genes (#pseudo genes), and the number of TALE classes (#classes).

| Technology | Assembly | Polishing | #TALEs | #iTALEs | #pseudo genes | #classes |
| --- | --- | --- | --- | --- | --- | --- |
| Illumina | MaSuRCA | none | 1 | 0 | 1 | 0 |
| PacBio | HGAP4 | none | 13 | 1 | 9 | 13 |
| PacBio | HGAP4 | Illumina (pilon) | 15 | 1 | 7 | 13 |
| ONT | Flye | comp. correction | 20 | 1 | 0 | 19 |
| ONT | Flye | Illumina (pilon) | 19 | 1 | 1 | 18 |
| ONT | Flye | PacBio (Arrow) | 20 | 1 | 0 | 19 |
| ONT | Flye | Homopolish (bacteria) | 17 | 1 | 1 | 16 |
| ONT | Flye | Homopolish (*X. oryzae*) | 17 | 1 | 1 | 16 |
| PacBio + Illumina | MaSuRCA | none | 0 | 0 | 4 | 0 |
| ONT + Illumina | Unicycler | none | 0 | 0 | 21 | 0 |

**Table 4.** Quality of assemblies based on sub-samples of fractions of the complete set of ONT reads. In all cases, the assembly yields contiguous sequences of the chromosome and two plasmids. u.u.: uncorrected, unpolished; i.p. Illumina-polished; c.c.: computationally corrected.

| Fraction | #reads | mean cov. | assembly | contig N50 | Busco | #TALEs | #iTALEs | #pseudo gene |
| --- | --- | --- | --- | --- | --- | --- | --- | --- |
| 100,00 % | 579,937 | 630 | u.u. | 4,989,365 | C:94.6% | 0 | 0 | 20 |
|  |  |  | i.p. | 4,989,976 | C:98.8% | 19 | 1 | 1 |
|  |  |  | c.c. | 4,989,458 |  | 20 | 1 | 0 |
| 75,00% | 434,953 | 473 | u.u. | 4,989,461 | C:93.8% | 0 | 0 | 20 |
|  |  |  | i.p. | 4,989,963 | C:98.9% | 20 | 1 | 0 |
|  |  |  | c.c. | 4,989,546 |  | 20 | 1 | 0 |
| 50,00% | 289,968 | 315 | u.u. | 4,989,470 | C:93.2% | 0 | 0 | 20 |
|  |  |  | i.p. | 4,989,960 | C:98.9% | 20 | 1 | 0 |
|  |  |  | c.c. | 4,989,557 |  | 20 | 1 | 0 |
| 25,00% | 144,984 | 159 | u.u. | 4,989,306 | C:92.2% | 0 | 0 | 19 |
|  |  |  | i.p. | 4,989,957 | C:98.9% | 20 | 1 | 0 |
|  |  |  | c.c. | 4,989,394 |  | 20 | 1 | 0 |
| 12,50% | 72,492 | 79 | u.u. | 4,989,133 | C:89.8% | 0 | 0 | 19 |
|  |  |  | i.p. | 4,989,945 | C:98.9% | 16 | 1 | 2 |
|  |  |  | c.c. | 4,989,220 |  | 20 | 1 | 0 |
| 6,25% | 36,246 | 40 | u.u. | 4,988,747 | C:85.2% | 0 | 1 | 17 |
|  |  |  | i.p. | 4,989,905 | C:98.8% | 13 | 1 | 3 |
|  |  |  | c.c. | 4,988,857 |  | 20 | 1 | 0 |
| 3,12% | 18,123 | 20 | u.u. | 4,988,237 | C:73.9% | 0 | 0 | 17 |
|  |  |  | i.p. | 4,989,885 | C:98.8% | 10 | 1 | 7 |
|  |  |  | c.c. | 4,988,371 |  | 20 | 1 | 0 |

**Table 5.** Comparison of the ONT-based assemblies of the three *Xoo* and the two *Xoc* strains on the level of TALEs. Per assembly, we record the total number of TALE genes (#TALEs), the number of iTALEs with a truncated C-terminus (#iTALEs), the number of TALE pseudo genes (#pseudo genes), and the number of TALE classes (#classes).

| Strain | #contigs | contig N50 | Busco | #TALEs | #iTALEs | #pseudo genes | #classes | #new classes |
| --- | --- | --- | --- | --- | --- | --- | --- | --- |
| *Xoo* PXO35 | 3 | 4,989,458 | 94.6% | 20 | 1 | 0 | 19 | 1 |
| *Xoo* FXO38 | 1 | 5,032,653 | 93.8% | 17 | 2 | 0 | 15 | 1 |
| *Xoo* Huang604 | 1 | 4,947,629 | 95.1% | 16 | 1 | 0 | 17 | 0 |
| *Xoc* BAI35 | 1 | 5,155,624 | 93.2% | 20 | 1 | 0 | 20 | 0 |
| *Xoc* MAI23 | 1 | 5,042,222 | 96.0% | 20 | 1 | 1 | 20 | 0 |

## RESULTS/DISCUSSION

### *Xoo* PXO35 – A genome with highly repetitive TALE genes only fully assembled from ONT reads

For *Xoo* PXO35, we generated reads from three alternative sequencing technologies, namely Illumina short reads, and PacBio and ONT long reads, each with their specific limitations and error profiles. This gives us the opportunity to systematically assess the assembly quality and contiguity achieved based on each of the technologies, or using hybrid and polishing approaches applied to combinations of these. Here, we especially focus on the correct assembly of TALE genes within their genomic context. TALE genes are notoriously difficult to sequence because of the repetitive structure of their DNA-binding domain (Fig. 2), which is composed of typically 34 amino acid long tandem repeats, corresponding to 102 bp on the DNA level. In addition, the DNA sequence coding for the N-terminal and C-terminal domains is highly conserved among TALE genes as well. Within the genomic sequence, TALEs are often organized in clusters of several TALE genes separated by short linker sequences (18), which further complicates a contiguous assembly, even from longer reads.

**Fig. 2.**
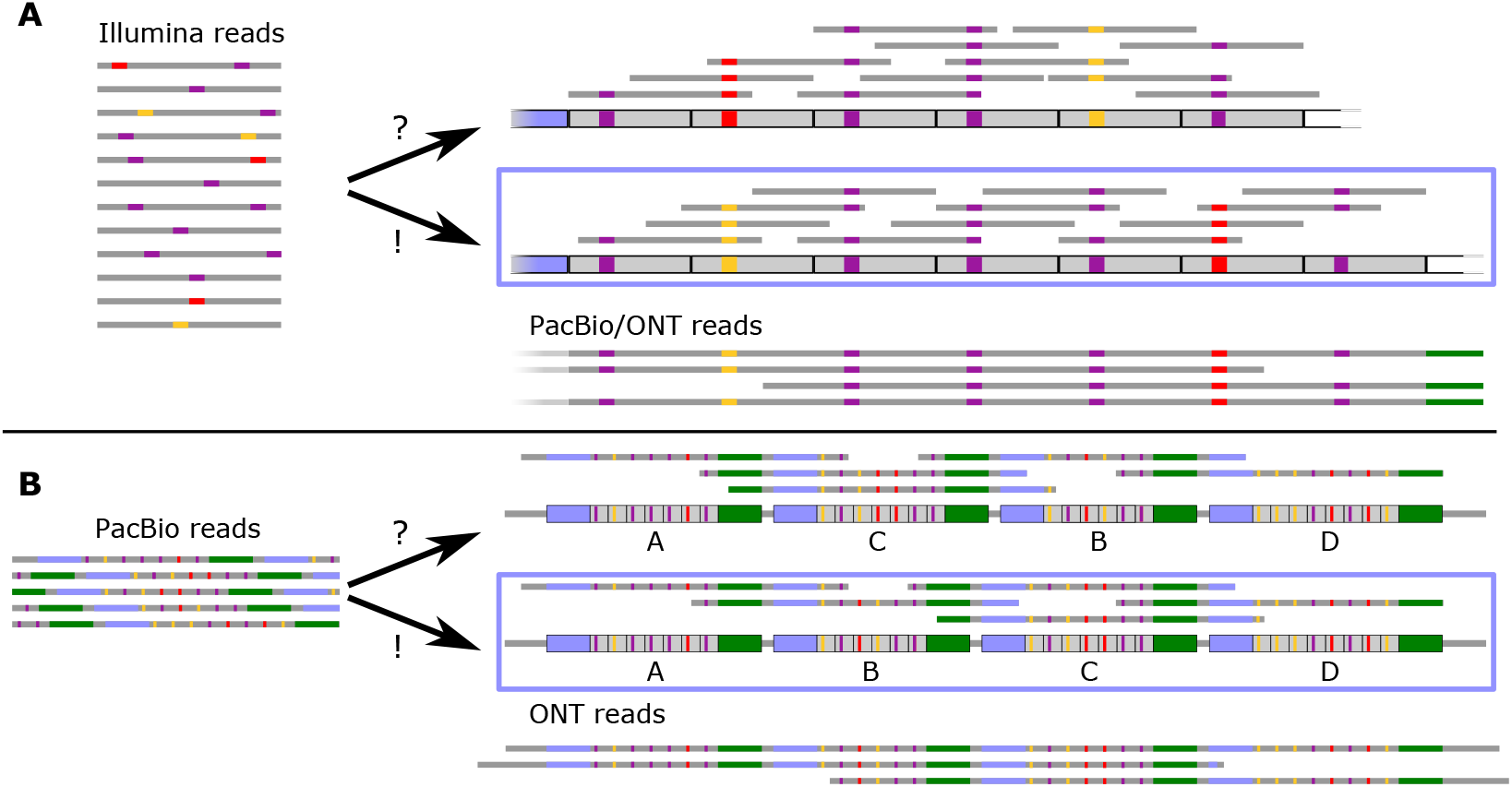
Schematic illustration of assembly problems based on read length. (A) Using short Illumina reads, the assembly of a single TALE may be ambiguous due to highly conserved tandem repeats, which often differ only in their RVD di-codon indicated by coloured boxes. Starting from the same set of Illumina reads, two alternative assemblies are consistent with the reads. Using long read technologies, however, the bottom assembly may be identified as the correct one. (B) Even using longer reads spanning complete TALEs, the organization and order of TALE within clusters may be ambiguous due to highly conserved N-terminus, C-terminus and linker sequence between TALEs. Here, very long ONT reads spanning complete TALE clusters are capable of resolving the arrangement of TALEs in large clusters.

In the following, we first compare assemblies based on reads from individual sequencing technologies and combinations thereof using general statistics of assembly contiguity and completeness. We then compare the assembly on a broader scale using genomic alignments with a focus on the placement and completeness of TALE clusters. Finally, we compare the TALE annotations on these assemblies with regard to the number of reconstructed TALEs and their assignment into TALE classes.

#### Comparison of assemblies by general statistics

For the purpose of comparing the different basic sequencing methods, we first assembled the genome of *Xoo* PXO35 in a non-hybrid manner from either only Illumina, only PacBio, or only ONT reads. For the assembly from Illumina reads only, we used the MaSuRCA method which is a hybrid approach that allows variable read lengths and significant sequencing errors (48). For the assembly based solely on short Illumina reads, we obtain a very large number of contigs (314) with a relative low N50 weighted median contig length of 26,708 bp. To assess the assembly with regard to the genes they contain, we used BUSCO (54), which relies on single-copy orthologs to quantify completeness. Applying the 1,152 BUSCO markers of the “xanthomonadales_odb10” dataset, the Illumina assembly still had a decent Busco score indicating that despite the low contiguity, many genes are still fully covered by a contig (cf. Table 2).

PacBio reads were de novo assembled using the classical hierarchical genome-assembly process 4 (HGAP4) pipeline (SMRT LINK v.7.0.1) (49). Even the much longer PacBio reads did not succeed in closing a complete chromosome, resulting in five contigs. Two of these correspond to two plasmid sequences whereas the chromosomal sequence is fragmented into three contigs.

Here, contig N50 is substantially larger than for the Illumina-based assembly (4,203,214 bp) and a large fraction of the approx. 5 Mbp chromosome is covered by the largest contig. Busco completeness is on the same level as observed for the Illumina-based assembly. Polishing the PacBio-based assembly with Illumina reads using pilon only mildly affects contig N50 and leads to a small increase in Busco completeness.

The assembly applying Flye to the ONT reads results in a completely contiguous genome of *Xoo* PXO35 comprising one chromosomal and two plasmid contigs. One plasmid contig has high similarity to a plasmid of *Xoo* CIAT and *Xoo* BXO1 and the other plasmid contig has high similarity to a plasmid of *Xoo* IX-280. The contig N50 corresponds to the length of the complete chromosomal sequence. Due to the specific error profile of ONT reads resulting in a substantial number of insertions and deletions, Busco completeness is clearly lower than for the Illumina-based and PacBio-based assemblies. Since the computational correction proposed in this paper only affects the TALE genes, which are not covered by the Busco gene set, it does not influence Busco completeness. By contrast, polishing this assembly with pilon using Illumina reads recovers Busco completeness to a similar range as for the previous two sequencing technologies. Similar results are also obtained by polishing with Arrow using PacBio reads, and by polishing with Homopolish using either bacteria or *X. oryzae* as the reference.

The hybrid assembly by MaSuRCA jointly considering PacBio and Illumina reads leads to a larger number of contigs and a lower Busco completeness than using only PacBio reads in the HGAP4 pipeline. The hybrid assembly by Unicycler jointly considering ONT and Illumina reads, however, yields a similar quality of the assembly as the ONT-based assembly with Illumina polishing.

In summary, we conclude that ONT reads are essential to assemble a fully contiguous *Xoo* PXO35 genome. Combined with Illumina short reads, either in a polishing step or in a hybrid assembly approach, this also yields a competitive Busco completeness compared with the alternative options.

#### Comparison of assemblies by genomic alignments

We further compare the different assemblies by genomic alignments using progressiveMauve to obtain an overview of their concordance in the arrangement of larger genomic regions. We find that – as expected – all assemblies using ONT reads are in large agreement (Fig. 3). Here, the different polishing or assembly strategies lead to local differences in the assembled sequence, whereas genomic blocks (indicated by colour) stay in the same order and strand orientation. The general placement of TALE clusters is also identical between all four assemblies. The two PacBio-based assemblies (with and without pilon polishing) may be aligned well to the ONT assemblies, but two contig borders (at approx. 1,230 kbp and approx. 2,000 kbp) are located within TALE clusters according to the ONT-based assemblies. By contrast, for the hybrid assembly based on PacBio and Illumina reads we may visually perceive the large number of contigs with frequently changing strand orientations in the alignment. Again, many of the contig borders are placed within TALE clusters. The solely Illumina-based assembly, finally, results in a highly fragmented assembly, which is lacking the majority of TALE clusters. We provide an archive containing the progressive-Mauve alignment file for further exploration as Supplementary File S1.

**Fig. 3.**
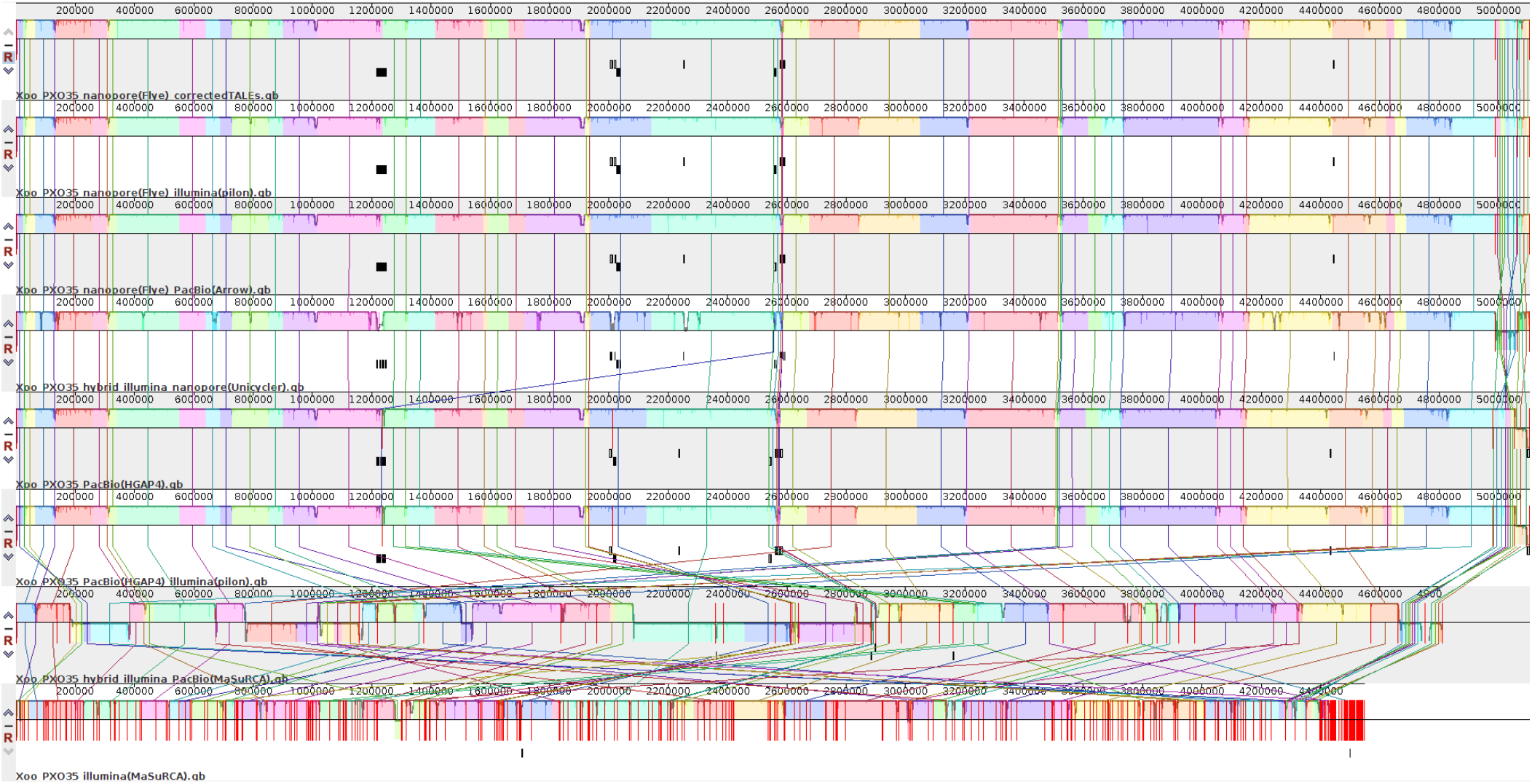
Complete alignment of different PXO35 assemblies using progressiveMauve. Contig borders are marked by red vertical lines. All ONT-based assemblies yield one contiguous chromosome and two plasmid sequences, whereas PacBio-based assemblies are split into multiple, long contigs with TALE clusters located at the contig borders. The hybrid and Illumina-based assemblies are characterized by a large number of smaller contigs. The location of TALE genes is indicated by boxes below the asssembly representation.

The genomic alignments of the different assemblies show that major differences between the ONT-based and the PacBio-based assemblies (and also the Illumina-based assembly) lie within large TALE clusters (Supplementary Fig. S2). To gain further insights into these differences, we consider the region around the first contig border within the PacBio-based assembly in Fig. 4. From the ONT-based assembly, we observe that this cluster contains seven TALEs from different TALE classes. The PacBio-based assembly correctly reconstructed the first two (BJ, AE) and the last two (AQ, AP) of these TALEs, while the remaining TALEs are located on either border of the two contigs and fail to be annotated and classified by AnnoTALE. Finally, the complete TALE cluster is missing from the Illumina-based assembly, where the end of a first contig is placed upstream, while the start of the following contig is located already downstream of this TALE cluster. However, it might be still questionable if the ONT-based assembly of this TALE cluster is indeed correct. Hence, we further consider ONT and PacBio reads mapped to the ONT-based assembly in Supplementary Fig. S3. We find that a large number of long ONT reads spans the complete TALE cluster, which provides further confidence in this assembly result. PacBio reads map also well to this region, although these had not been sufficient to reconstruct this TALE cluster initially. In general, each of the TALE clusters or single TALEs within the genome is fully covered by a sufficient number of ONT reads, which increases confidence in their correct assembly (Supplementary Table C).The second contig border within the PacBio-based assembly (at approx. 2,000 kbp) is also located within a large TALE cluster and we further consider it in Supplementary Fig. S4. The cluster observed in the ONT assembly contains TalAB, TalAL and TalBA on the forward strand as well as TalAC, TalAS and TalAG on the reverse strand. However, in the PacBio-only assembly the TALEs on the forward strand are TalAB, a truncated TalAL and on the reverse strand AS and AG, which means that the two TALEs TalBA and TalAC in the center of the large cluster could not be assembled correctly. In order to further investigate the correctness of this ONT-based TALE cluster, we have considered the mapped ONT and PacBio reads in Supplementary Fig. S5, and find that this region is well covered by long ONT reads as well as the shorter PacBio reads. In summary, our results indicate that both long-read techniques are generally suited to assemble *Xanthomonas* genomes containing (clusters of) TALE genes as shown previously (4, 60). However, in case of very large TALE clusters, as in the first and second example with a cluster of seven and six TALEs, respectively, spanning 35 kbp, the repetitive nature and high conservation of TALE genes may require typically longer ONT reads for a complete assembly result.

**Fig. 4.**
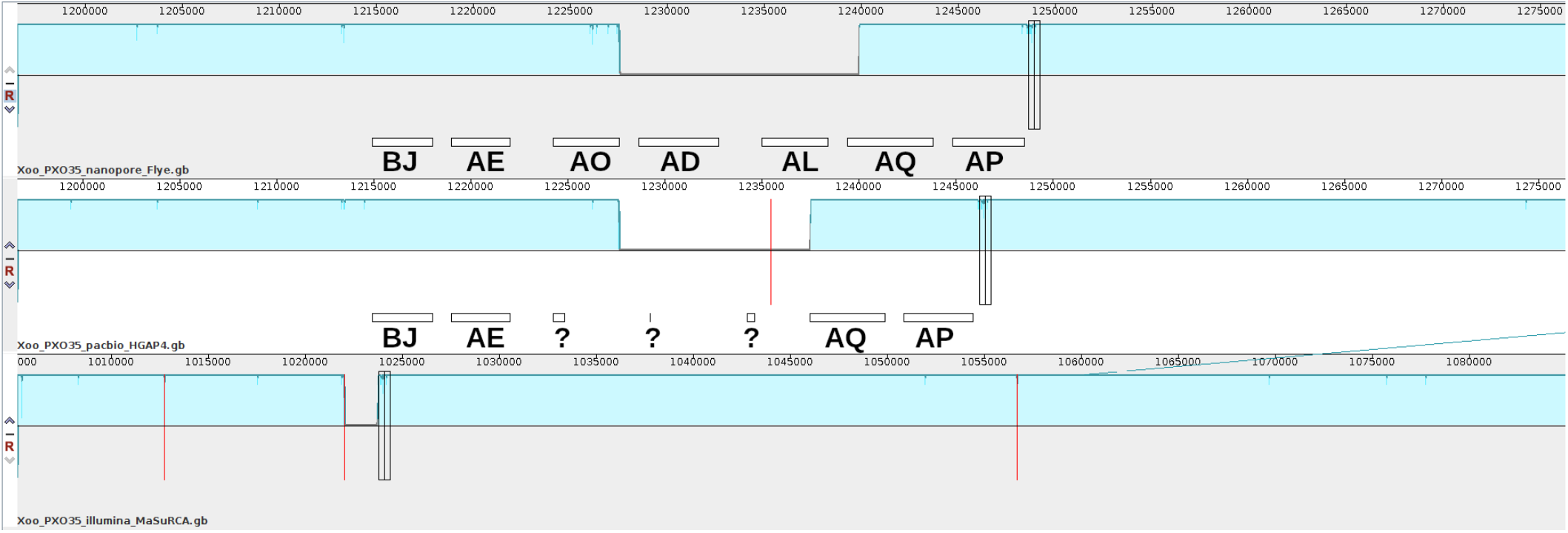
A large cluster of repetitive TALE genes in *Xoo* PXO35 at approx. 1,220 kbp prevents a contiguous assembly. Contig borders are marked by red vertical lines. In the ONT-based assembly, seven TALEs are present in this cluster (boxes), whereas only four of them could be reconstructed from the PacBio-based assembly and none in the Illumina-based assembly. The black vertical rectangles represent the homologous position between the three assemblies.

ONT sequencing entails specific problems with regard to insertions and deletions, especially in homopolymer regions (61). Such sequencing errors may also be observed within TALE genes, which are in the focus of this manuscript. With the assemblies of the *Xoo* PXO35 chromosome polished by either Illumina or PacBio reads, we get insights into the prevalence of these sequencing errors and also derive and test a computational method for their correction in ONT-only assemblies.

In Fig. 5, we present a cartoon of the genomic sequence of one TALE (TalBJ8) from *Xoo* PXO35. Here, we assume that the Illumina-polished and PacBio-polished variants of this genomic region have the correct base sequence. In addition, this TALE is located in the portion of the large TALE cluster (cf. Fig. 4) that could be correctly assembled from PacBio reads alone. Hence, we have three reference sequences against which we may compare the sequence of the ONT-based assembly. Here, we find four homopolymer stretches, two in the sequence coding for the N-terminus and two in the sequence coding for the C-terminus that appear to be affected by deletions in the ONT-based assembly. Notably, these deletions occur despite the very high coverage with ONT reads (∼662 X), and are unlikely to be resolved by even deeper sequencing.

**Fig. 5.**
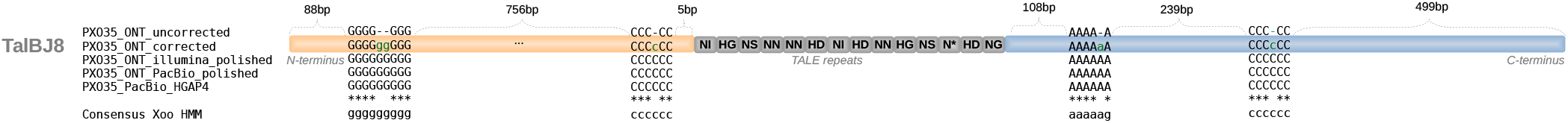
Typical errors in homopolymers for ONT sequencing and how these may be corrected. The initial ONT-based assembly is lacking nucleotides in homopolymer stretches, which may be corrected computationally by comparison to the corresponding consensus from the HMM. The resulting corrected sequence is in accordance with the genome version after Illumina-based or PacBio-based polishing of the initial ONT-based assembly.

Here, we propose a computational correction method for ONT-only assemblies based on hidden Markov models learned from the sequences of N-terminus, C-terminus, and individual repeats of known *Xoo* or *Xoc* TALEs. In all four cases, we observe that the consensus of the corresponding HMM as aligned using nhmmer matches the sequence of the three reference sequences at the deletion sites. As such errors predominantly appear within homopolymers, we correct these by filling the deletion sites with the homopolymer nucleotide. In this case, this procedure yields the same result as simply transferring the consensus nucleotide of the HMM to a corrected version of the ONT-based sequence. Outside of homopolymer stretches we, hence, use the HMM consensus for computational correction. Optionally, this may be followed by a read-based custom substitution polishing, which is based on the ONT reads mapped back to the corrected sequence. If in those mapped reads another nucleotide appears at least 1.5 times as frequent as the previous correction choice, the nucleotide from the mapped reads is used instead. With the computational correction at hand, we may now reconstruct complete TALomes of *Xanthomonas* strains based on ONT reads only without the need for additional Illumina or PacBio reads used for polishing. However, this comes at the potential risk that true deletions within TALEs are corrected by mistake. Hence, we indicate the sites of computational corrections by lower case letters to make them identifiable in downstream analyses.

#### Comparison of annotated TALEs

In the remainder of this section, we compare the different assemblies, including the ONT-only assembly combined with the computational correction of TALE genes, on the level of their TALomes as annotated by AnnoTALE. We find that – as already indicated by the genomic alignments – the Illumina-based assembly is able to reconstruct a single TALE gene and one further TALE pseudo gene only (Table 3). The PacBio-based assemblies with and without pilon polishing yield a substantially larger number of TALE genes, but contain a larger number of pseudo genes than might be expected. Still, polishing by pilon helps to recover two of these pseudo genes. A subset of pseudo genes is located within the large TALE cluster as indicated by the question marks in Fig. 4. The largest number of TALE genes is annotated from the ONT-based assemblies. Here, we find up to 20 TALE genes and one iTALE (55, 56). iTALEs are truncated versions of TALEs which don’t act as gene activators, but instead block TALE recognition by specific rice resistance proteins (55, 56). They are therefore functionally different from normal TALEs. Notably, the ONT-only assembly combined with the computational correction yields the same TALEs, iTALE, and TALE classes as the PacBio-polished variant. The hybrid assembly based on PacBio and Illumina reads, which already showed low contig N50 and Busco completeness in the general assessment, does not contain a single complete TALE gene and only four TALE pseudo genes. Surprisingly, also the ONT + Illumina hybrid assembly that appeared to be of high quality in the general assessment contains only TALE pseudo genes, although the number of pseudo genes indicates that the total set of TALE regions might be complete. One reason for this observation could be that the internal correction of ONT reads using Illumina reads within the Unicycler pipeline fails to adequately correct the highly conserved N-terminus and C-terminus of TALE genes, but especially the highly conserved tandem repeats including the di-codons of RVDs.

Typically, ONT sequencing suffers from problems within homopolymers. Homopolish may improve assemblies in homopolymeric regions based on homologous sequences. Applying Homopolish to the ONT-based assembly of *Xoo* PXO35 results in a reduced number of annotated TALEs compared with the other polishing/correction options due to the introduction of frame shifts and/or premature stop codons. More importantly, we observe that Homopolish may introduce modifications in the di-codon of TALE repeats and, subsequently, their RVD sequence. In the case of *Xoo* PXO35, this issue occurs for both Homopolish reference data sets, i.e., bacteria and *X. oryzae*. Supplementary Table B lists all changes within the RVDs of all TALEs of PXO35 for both Homopolish variants, while Supplementary Fig. S6 shows the effect of Homopolish in comparison to other polishing variants within the RVD sequence of TalAO. As polishing with Homopolish may lead to altered RVD sequences, we consider it unsuitable for polishing TALE genes, since a correct RVD sequence is essential for downstream analyses such as the prediction of TALE targets.

In summary, the comparison of different assembly, polishing, and correction strategies leads to the following conclusion for sequencing TALomes of *Xanthomonas* bacteria, which may likely also apply to similar repetitive structures within genomes: First, long ONT reads are required to yield fully contiguous genomes of strains with large TALE clusters. Second, further polishing by Illumina or PacBio reads is necessary if one is interested in a complete gene annotation. Using Homopolish as an alternative may recover many protein coding genes, but may also introduce undesirable modifications within TALE genes. Third, hybrid assembly strategies starting from an initial short-read graph may fail for conserved, repetitive regions like TALEs. Fourth, ONT-only assemblies combined with a computational correction step are sufficient to yield complete and correct TALomes of *Xanthomonas* bacteria. Hence, we follow the latter strategy in the remainder of this manuscript and focus on TALE genes.

### Multiplexing capacity of complete TALome-decoding from *Xanthomonas* strains using ONT sequencing

As mentioned previously, the total set of ONT reads yields a very high coverage of the approx. 5 Mbp genome of *Xoo* PXO35, although the true mean coverage after re-mapping the reads to the assembly is slightly lower than could be expected from the raw reads (cf. Table 1). However, it remains an open question how much coverage is indeed required to correctly reconstruct a TALome from ONT reads or, put differently, how many genomes of TALE-carrying *Xanthomonas* strains could be multiplexed to be sequenced on a single ONT flow cell. The rather complex organization of the *Xoo* PXO35 genome with its large TALE clusters and two plasmids combined with a large number of available ONT reads provides an ideal scenario to test the potential multiplexing capacity *in silico*.

To this end, we perform the assembly using Flye on sub-samples of different fractions of the original ONT reads, followed by Illumina-based polishing or our computational correction, respectively.

We find that the contig N50 of the resulting assemblies stays highly stable across the whole range of sub-sampling fractions (Table 4). Even from a subset of only 3.12% of the original ONT reads, Flye is able to generate a fully contiguous assembly of the chromosome and the two plasmids. Without any polishing, the Busco completeness starts to drop early, and only yields 73.9% of the Busco genes for the lowest number of reads. However, when polishing with Illumina reads, Busco completeness stays rather constant as well. As the computational correction does only affect TALE genes, Busco completeness after computational correction is identical to the uncorrected, unpolished case.

Regarding the reconstruction of TALE genes, we find that without polishing or correction, none of the TALE genes is reconstructed correctly leading to a large number of pseudo genes due to insertions or deletions. Illumina polishing of the ONT-based assembly yields an (almost) complete set of TALEs and iTALE down to a fraction of 25% of the original ONT reads. With even lower sub-sampling fractions, however, the number of correct TALEs drops, which is only partly explained by these TALE becoming pseudo genes. By contrast, the computational correction is able to correctly reconstruct all TALE genes and the iTALE even for the lowest sub-sampling fraction. Hence, it would have been possible to determine the complete TALome of *Xoo* PXO35 from less than 20, 000 ONT reads, which illustrates the huge multiplexing capacity of ONT sequencing combined with a computational correction of TALE genes.

### TALEs and TALE classes present in strains

Having established a pipeline for assembling fully contiguous *Xanthomonas* genomes from ONT reads combined with a computational correction of TALE genes, we applied this pipeline to two further *Xoo* strains, namely FXO38 and Huang604, and to two African *Xoc* strains, BAI35 and MAI23.

For these remaining strains with ONT data generated in this study, each assembly leads to a single, circular chromosome with a length in the expected range of approx. 5 Mbp (Table 5). Busco completeness is in the same range as observed for *Xoo* PXO35 previously. Here, we observe the lowest Busco completeness for *Xoc* BAI35 (93.2%) and the highest Busco completeness for *Xoc* MAI23 (96.0%). The number of TALEs annotated for these genomes using AnnoTALE after computational correction of TALE genes varies between 16 (*Xoo* Huang604) and 20 (*Xoo* PXO35, *Xoc* BAI35 and MAI23), with up to one additional pseudo gene, and one to two iTALEs per strain.

From the genomic alignments of the assemblies of the three *Xoo* strains (Fig. 6A), we observe substantial rearrangements and inversions, which also affect the location and composition of TALE clusters. Compared with *Xoo* PXO35, TALE clusters of the other two *Xoo* strains contain a lower number of TALEs per cluster, where the largest cluster is composed of five (FXO38) and four (Huang604) TALE genes. The TALEs of the large cluster in *Xoo* PXO35 (TalBJ–TalAP) either do not occur in FXO38 and Huang604 (TalBJ, TalAL), are part of smaller clusters (TalJQ–TalAD in FXO38 and TalAE– TalAP in Huang), or are located in another cluster together with an additional TALE (TalAL–TalAD in Huang604). In all three genomes, the iTALE TalAI is located outside of clusters. There are two types of iTALEs, type A and type B, that differ in the C-terminal truncation (55, 56, 62, 63). TalAI from PXO35 and Huang604 are type A iTALEs, whereas TalAI from FXO38 is a type B iTALE. In addition, FXO38 contains a type A iTALE with TalDQ.

**Fig. 6.**
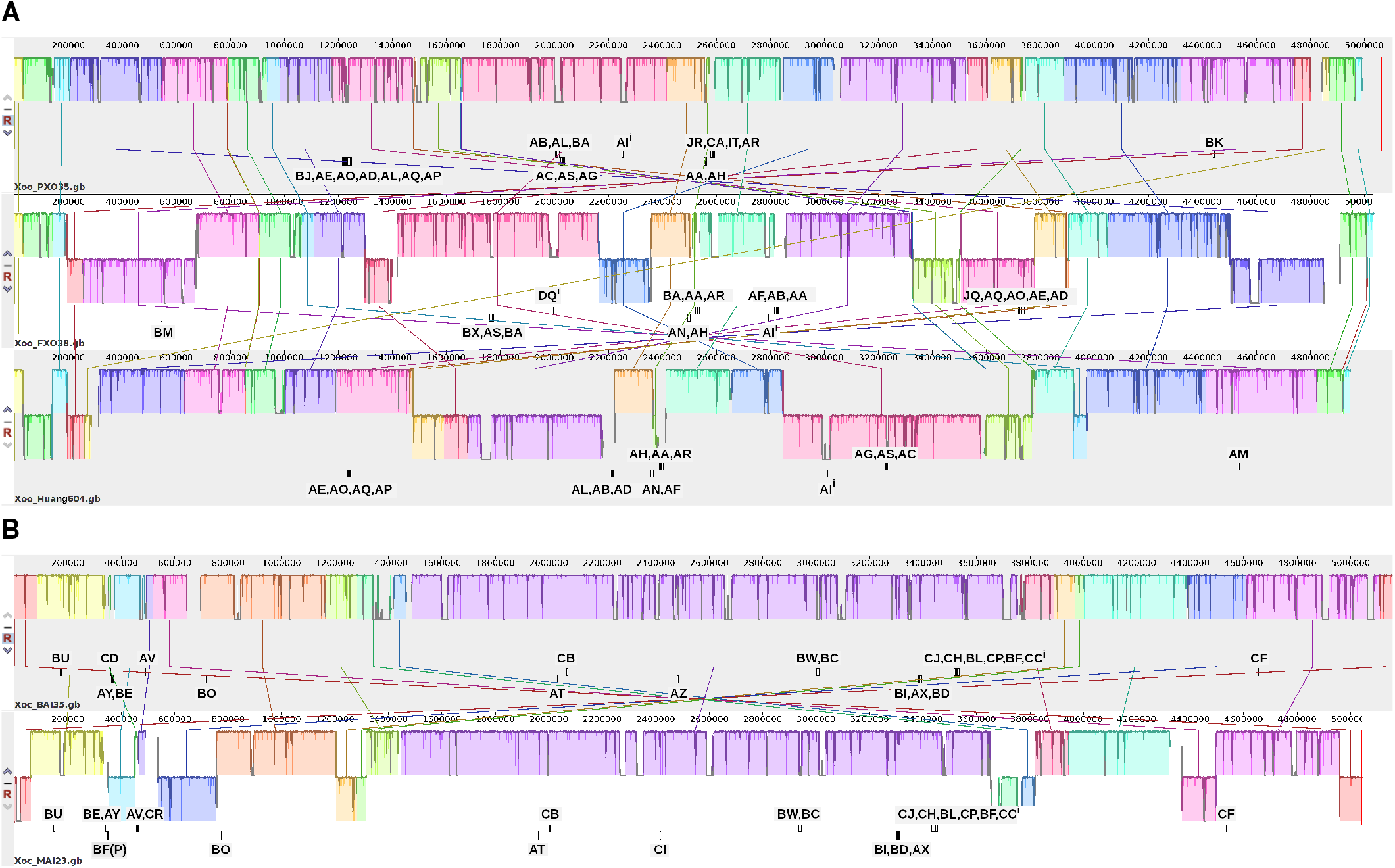
Complete alignment of assemblies using progressiveMauve. (A) Alignment of the three *Xoo* strains PXO35, FXO38 and Huang604. (B) Alignment of the two *Xoc* strains BAI35 and MAI23. Each iTALE is marked with a superscript i.

For the two *Xoc* strains (Fig. 6B), the number of rearrangements and inversions is lower with a large collinear block between approx. 1.5 Mbp and 3.7 Mbp. Within this block, also the location and composition of TALE clusters is largely conserved with one single TALE replaced (TalAZ vs. TalCI) and one switch of TALE positions within a cluster (TalBI, TalAX, TalBD vs. TaBI, TalBD, TalAX). In both strains, many TALEs are located outside of clusters as single TALEs and both have one iTALE of type B (TalCC).

We further compared the TALomes of the newly sequenced strains to those of well-known strains, namely *Xoo* PXO99^*A*^ and Asian *Xoc* BLS256 (Fig. 7). In general, we find a clear distinction between the TALE classes of the *Xoo* and *Xoc* strains. The two classes that may be present in both pathovars (TalAD and TalAF) do not occur in the two newly sequenced *Xoc* strains. TalAD is present in all three *Xoo* strains, while TalAF is missing in *Xoo* PXO35. Several further common *Xoo* TALEs are present in all three *Xoo* strains (TalAA, TalAB, TalAE, TalAH, TalAI, TalAO, TalAQ, TalAR, TalAS). *Xoo* PXO35 has many unique TALEs that are not present in the other strains (TalBJ, TalBK, TalCA) but have been observed in *Xoo* strains previously. By contrast, the TALome of Huang604 appears to be composed of only common TALEs.

**Fig. 7.**
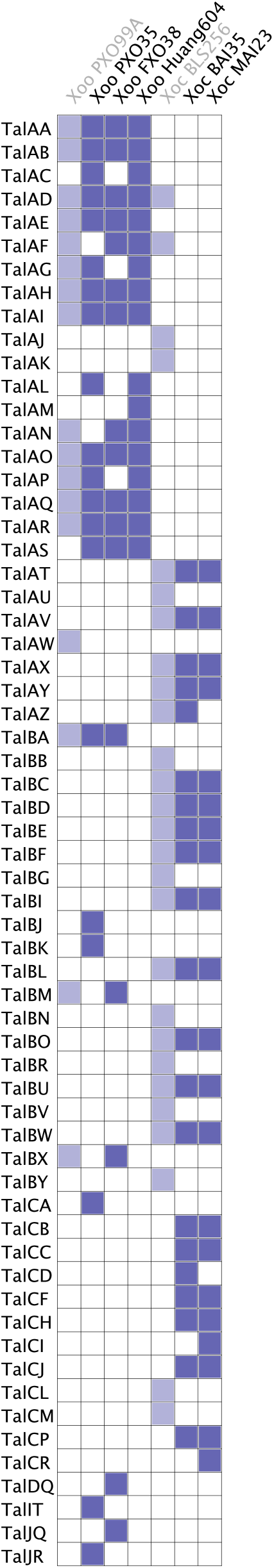
Presence of TALE classes in 5 ONT sequenced strains compared to 2 well-known strains. The nomenclature for TALEs is based on AnnoTALE (4). AnnoTALE groups similar TALEs according to their RVDs into classes, as indicated by the last two capital letters in their name. Accordingly, TALEs from the same class can be expected to target the same plant gene(s). To further distinguish TALEs from different strains within a class an allele number is added which is omitted here, because only TALE classes are shown.

For the two *Xoc* strains, the situation is similar with many common *Xoc* TALEs (TalAT, TalAV, TalAX, TalAY, TalAZ, TalBC, TalBD, TalBE, TalBF, TalBI, TalBL, TalBO, TalBU, TalBW) but also some less frequent TALEs (TalCB, TalCC, TalCF, TalCH, TalCJ, TalCP) that have been observed in other *Xoc* strains but are not present in BLS256. TalCD and TalAZ are present in BAI35 but not in MAI23, while TalCI and TalCR are present in MAI23 but not in BAI35. However, TalCD, TalAZ, TalCI and TalCR have all been observed in other *Xoc* strains before (4). A graphical overview of the TALEs in all fully sequenced *Xoo* and *Xoc* strains can be found in Supplementary Figures S7 and S8, respectively.

## Conclusions

We have solved the composition of the largest TALE cluster in *Xoo*, so far, which contains seven individual TALEs and is a unique rearrangement between cluster VIII and IX. *Xoo* strain PXO35 was originally isolated in 1972 in the Philippines (64) and belongs to race 1 which is characterized by a relative high diversity between strains. Although the virulence of this strain is high, this particular TALE cluster composition does not occur in other *Xoo* strains from this geographic location. It is possible, that this strain has a disadvantage on certain rice varieties in comparison to more frequently found strains. Alternatively, we might still underestimate the full diversity and frequency of TALE rearrangements in native *Xoo* populations.

ONT reads are perfectly suited to yield highly contiguous genomes of *Xanthomonas* bacterial strains that could not be resolved based on Illumina or PacBio reads due to clusters of highly repetitive *TALE* genes. While polishing by, e.g., Illumina reads, is still required to obtain complete and correct gene models of all bacterial genes, the computational correction pipeline presented in this manuscript is capable of correctly reconstructing the TALomes of *Xanthomonas* strains from ONT reads alone. As the quality of ONT reads is developing rapidly with new types of flow cells and improved chemistry, it can be expected that ONT sequencing will become the standard method for sequencing *Xanthomonas* strains soon. Due to the moderate coverage required for obtaining complete TALomes of *Xanthomonas* strains, sequencing on an ONT flongle might be sufficient for studying individual strains.

Given the huge multiplexing capacity of ONT sequencing combined with the computational correction of TALE genes, it is now possible to sequence large numbers of *Xanthomonas* strains in parallel to address pathogen populations and characterize strains from many more geographic regions. The expected expansion of TALE sequences is a good argument to support our unified Anno-TALE (http://www.jstacs.de/index.php/AnnoTALE) nomenclature, which groups similar TALEs into classes with a common name and allows for a quick understanding of TALE cluster compositions (Fig. 7, Supplementary Fig. S7, Supplementary Fig. S8) (4). The ONT pipeline presented here may lead to new insights into the importance and prevalence of certain TALE classes, but also evolutionary mechanisms shaping their TALomes.

## FUNDING

This work was supported by grants from the Deutsche Forschungsgemeinschaft (http://www.dfg.de) (BO 1496/8-2 to JB, GR 4587/1-2 to JG and FZT 118/2 to SK and MM). Work in RK’s laboratory was supported by a grant from the French Agence Nationale de la Recherche (ANR-2010-BLAN-1723). MM and SK were supported by the Ministry for Economics, Sciences and Digital Society of Thuringia (TMWWDG), under the framework of the Landesprogramm ProDigital (DigLeben-5575/10-9).

This preprint is formatted using a LATEX class by Ricardo Henriques

