## Supplementary Figures S1 - S8, Supplementary Tables A and B, Supplementary Methods for "Assembling highly repetitive *Xanthomonas* TALomes using Oxford Nanopore sequencing"

### SUPPLEMENTARY MATERIAL

#### Supplementary Figures

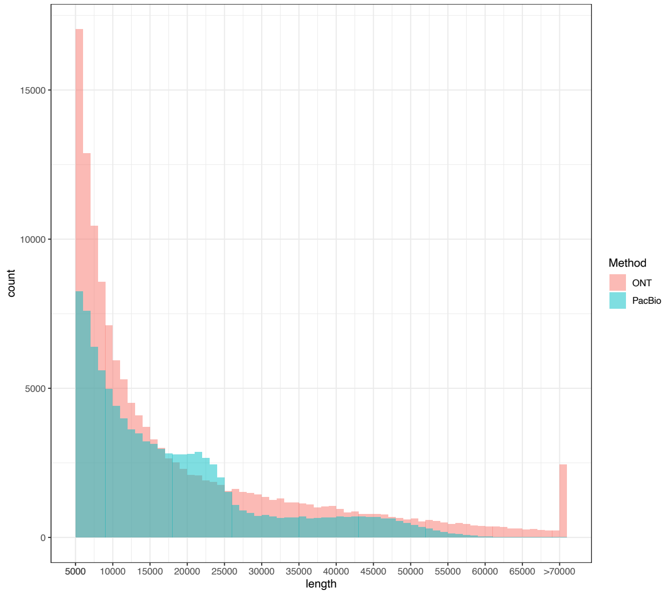


**Supplementary Figure S1.** Histograms of the read lengths above 5 kbp in PacBio and ONT libraries for *Xoo* PXO35. Read lengths above 70 kbp are aggregated in a single bin. A number of $45,582$ PacBio reads and $439,861$ ONT reads have lengths below 5 kbp.

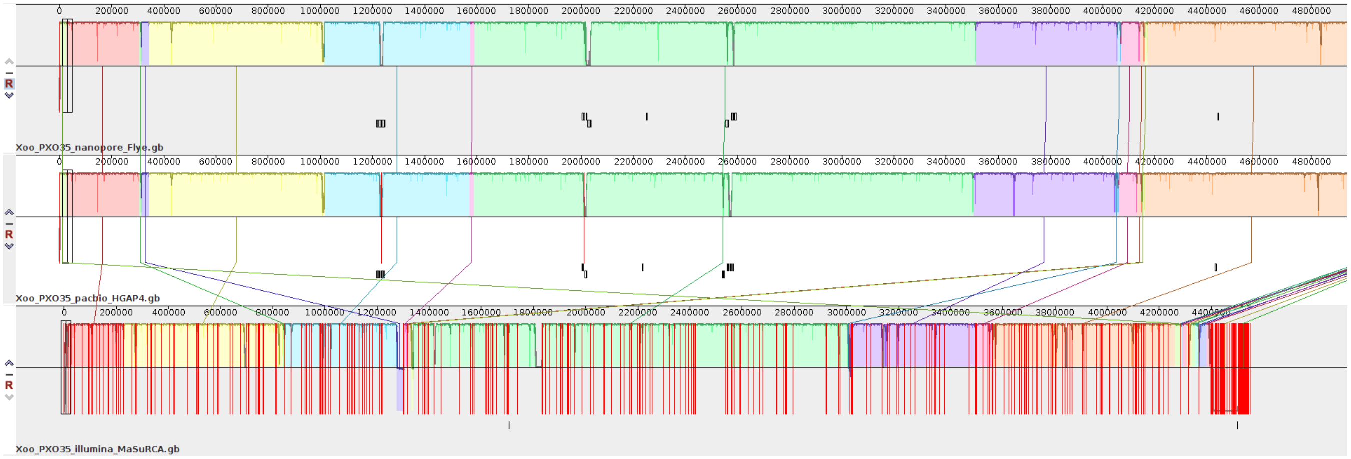


**Supplementary Figure S2.** Genomic alignment of ONT-based, PacBio-based and Illumina-based PXO35 assemblies using progressiveMauve for a subset of the assemblies shown in Figure 3. Contig borders are marked by red vertical lines. Large TALE clusters in the ONT-based assembly are located at contig borders of the PacBio-based assembly.

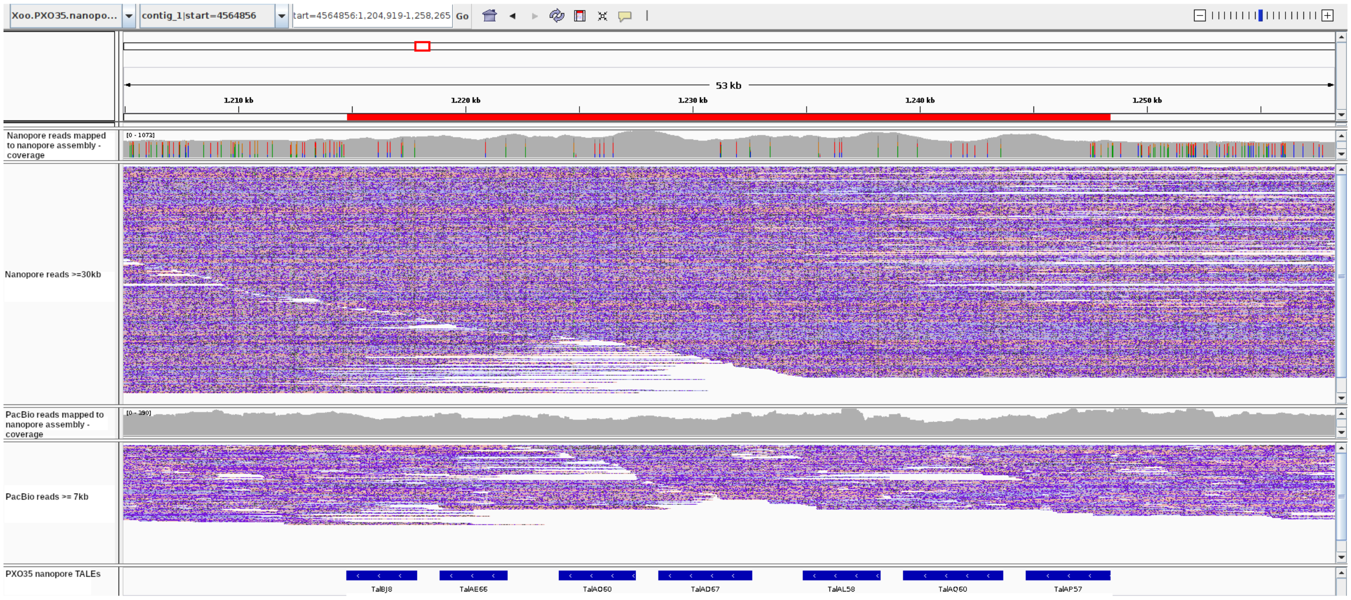


**Supplementary Figure S3.** IGV screenshot of the ONT-based, computationally corrected assembly of *Xoo* PXO35 in the region of a large TALE cluster at approx. 1,220 kbp. The first coverage track refers to all ONT reads that map to the region of the cluster. The corresponding alignment track contains only long ONT reads with a length of at least 30 kbp. The second coverage track refers to all mapped PacBio reads and the corresponding alignment track only includes PacBio reads with a length of at least 7 kbp. Mapped reads are sorted according to start location. The large TALE cluster reconstructed from the ONT reads using Flye is well-supported by ONT but also PacBio reads.

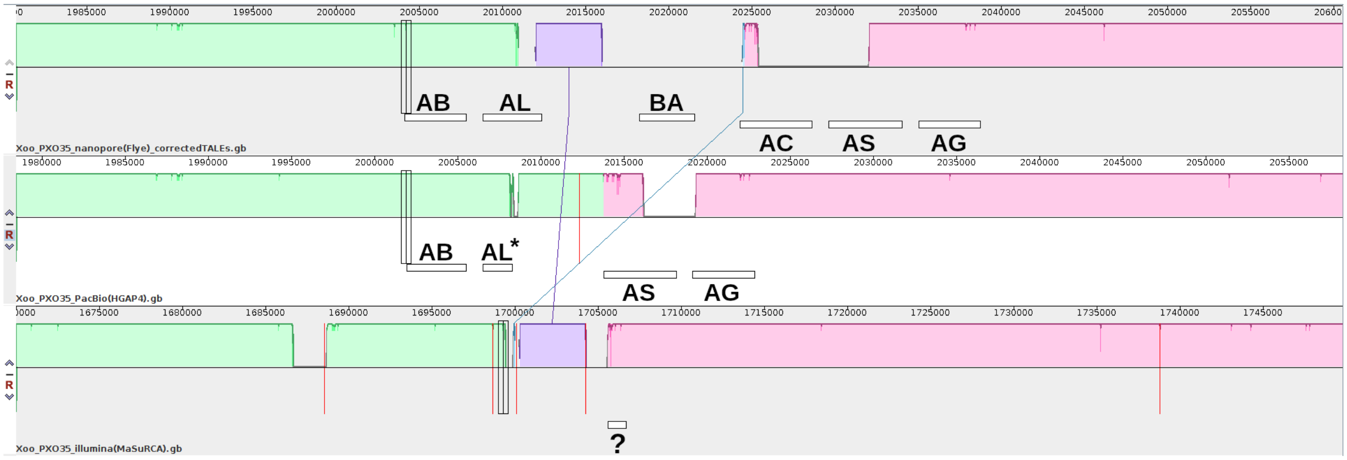


**Supplementary Figure S4.** A large cluster of repetitive TALE genes in *Xoo* PXO35 at approx. 2,000 kbp prevents a contiguous assembly. The asterisk indicates a truncated TALE gene. Contig borders are marked by red vertical lines. The large TALE cluster cannot be completely resolved from PacBio reads, while TALEs are missing in the Illumina-based assembly.

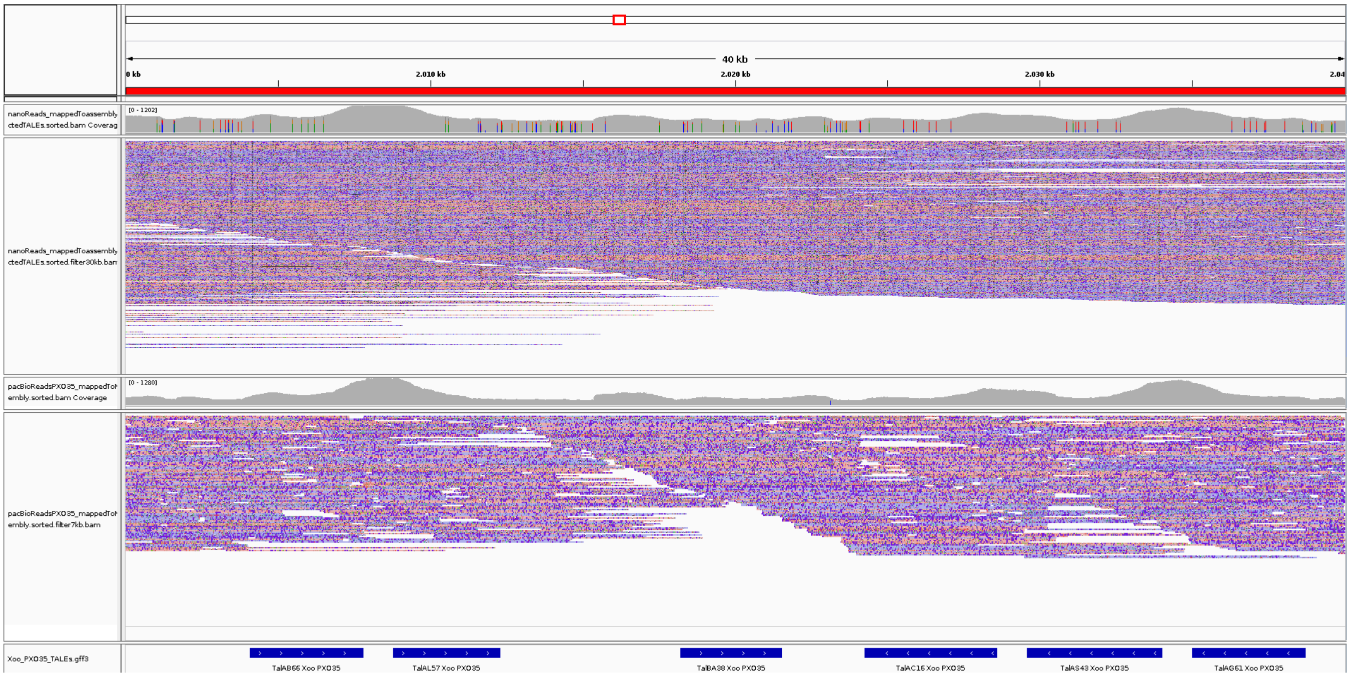


**Supplementary Figure S5.** IGV screenshot of the ONT-based, computationally corrected assembly of *Xoo* PXO35 in the region of a large TALE cluster at approx. 2,000 kbp. The first coverage track refers to all ONT reads that map to the region of the cluster. The corresponding alignment track contains only long ONT reads with a length of at least 30 kbp. The second coverage track refers to all mapped PacBio reads and the corresponding alignment track only includes PacBio reads with a length of at least 7 kbp. Mapped reads are sorted according to start location.

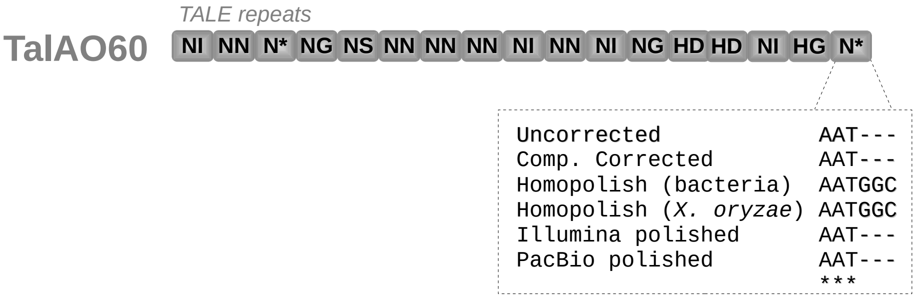


**Supplementary Figure S6.** Comparison of the polishing result of Homopolish using 'bacteria' or *X. oryzae* as a reference with the uncorrected assembly, the computationally corrected assembly, the assembly polished using Illumina reads, and the assembly using PacBio reads for TalAO60 of *Xoo* PXO35. We find that in both variants, Homopolish introduces a codon for an additional amino acid, which is not supported by the polishing variants based on experimental data (Illumina/PacBio).

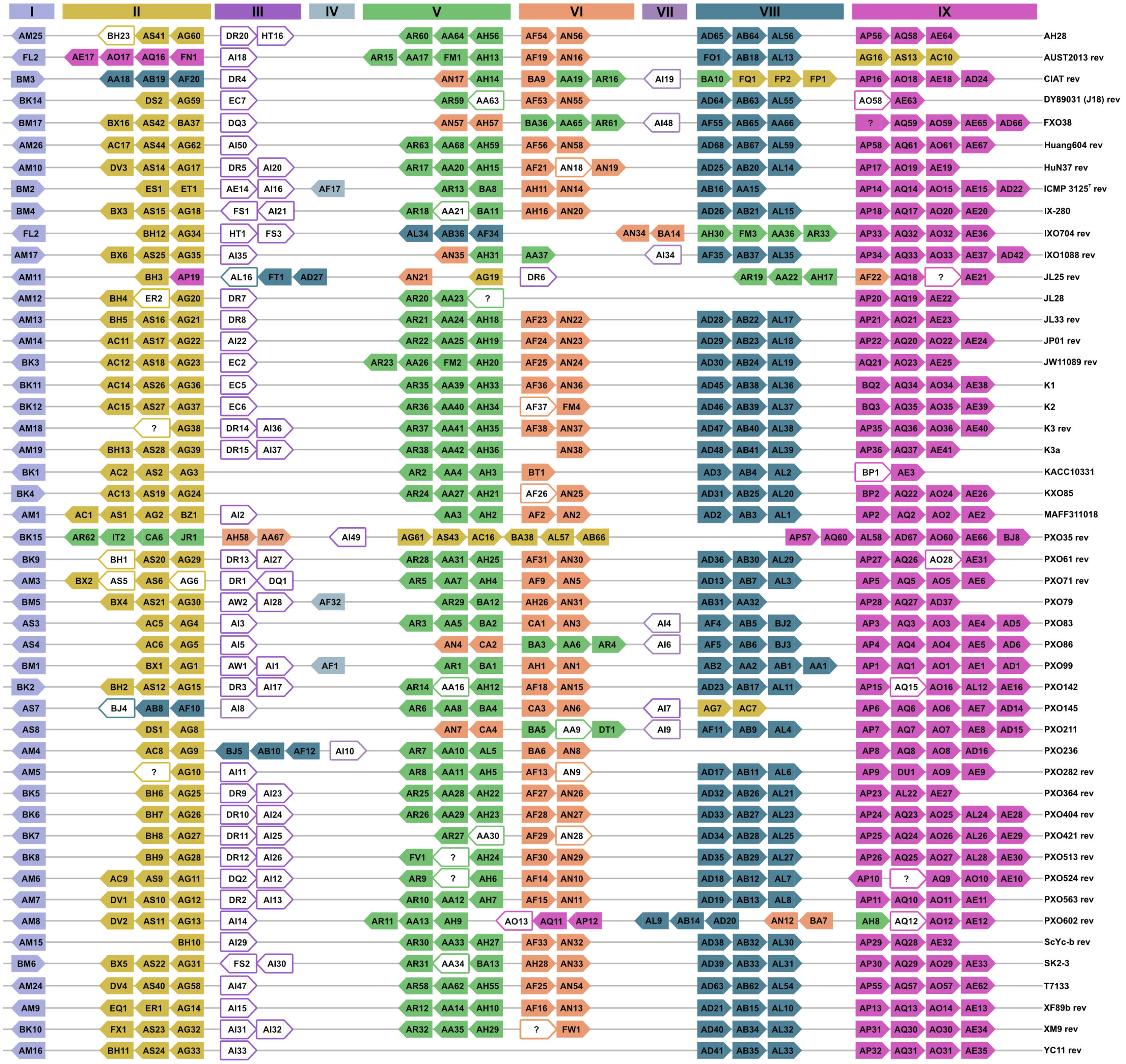

**Supplementary Figure S7.** Overview of TALE cluster assignment of sequenced Asian *Xoo* strains. TALEs are represented as arrows, directions indicate the relative orientation in the corresponding genome. All TALEs are assigned to classes using AnnoTALE and named accordingly. The two capital letters of the TALE class is shown and in addition an allele number which is unique for every particular TALE gene and assigned to distinguish TALEs from different bacterial strains. TALE clusters are defined at the top, affiliations are represented by colors. Strain names are shown at the right. Pseudo TALE genes are represented by a colored outlined white arrow. If a TALE could not be assigned to a class by AnnoTALE, the arrow is marked with a question mark. Strains with large genomic rearrangements may be shown reverse complemented for clarity, which is indicated by "rev" after the strain name.

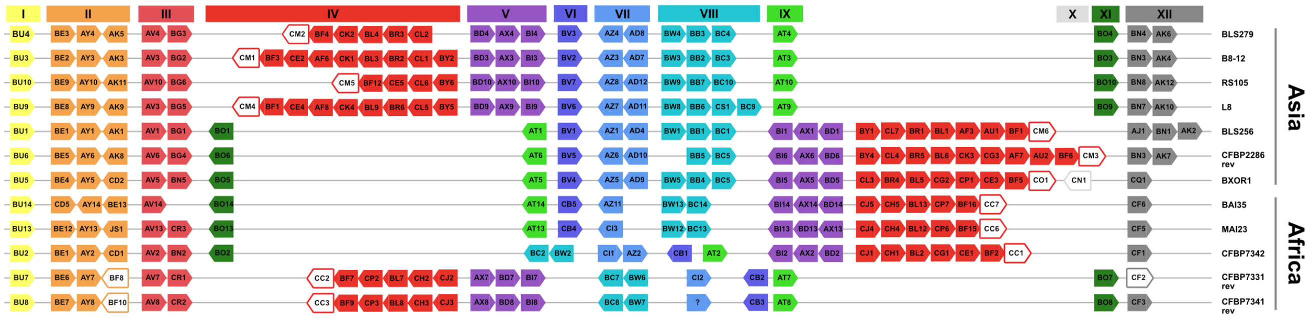


**Supplementary Figure S8.** Overview of TALE cluster assignment of sequenced Asian and African *Xoc* strains. TALEs are represented as arrows, directions indicate the relative orientation in the corresponding genome. All TALEs are assigned into classes using AnnoTALE and named accordingly. TALE clusters are defined at the top, affiliations are represented by colors. Strain names and geographical origin are shown at the right. Pseudo TALE genes are represented by a colored outlined white arrow. If a TALE could not be assigned to a class by AnnoTALE, the arrow is marked with a question mark. Strains with large genomic rearrangements are shown reverse complemented for clarity, which is indicated by "rev" after the strain name.

#### Supplementary Tables

**Supplementary Table A.** Statistics of read lengths in the PacBio and ONT library for *Xoo* PXO35.

| Read length | # PacBio reads | # ONT reads |
|---|---|---|
| $< 1000$ | $7,468$ | $252,612$ |
| $> 10000$ | $65,952$ | $84,042$ |
| $> 20000$ | $32,796$ | $46,786$ |
| $> 30000$ | $15,005$ | $29,486$ |
| $> 40000$ | $8,317$ | $17,932$ |
| $> 50000$ | $1,907$ | $10,257$ |
| $> 60000$ | $78$ | $5,338$ |
| $> 70000$ | $1$ | $2,447$ |

**Supplementary Table B.** Overview of the effect of Homopolish on annotated TALE genes in the *Xoo* PXO35 assembly. For each TALE that has been annotated after computational correction, we indicate (+/-) if the TALE is also present in the AnnoTALE annotation after polishing with Homopolish using 'bacteria' or *X. oryzae* as reference. Missing TALEs are the result of premature stops or frame shifts introduced by Homopolish within the TALE sequence. In addition, we list differences in RVDs introduced by Homopolish, which typically lead to the introduction of an additional, 13th amino acid in the repeat (e.g., N* → NG), or the deletion of the 13th amino acid (e.g., NN → N*). One specific example of such an introduction of a 13th amino acid is illustrated in Supplementary Figure S6. In some cases (TalAA, TalJR), the modifications introduced by Homopolish lead to a different class assignment of the corresponding TALE.

| PXO35 TALEs (comp. corrected) | Homopolish (bacteria) | RVD differences | Homopolish (*X. oryzae*) | RVD differences |
|---|---|---|---|---|
| TalAA | + | - | TalJT | HG → H* |
| TalAB | + | - | + | - |
| TalAC | - | | + | - |
| TalAD | + | - | + | - |
| TalAE | + | - | + | - |
| TalAG | + | - | + | - |
| TalAH | + | N* → NG | + | N* → NG |
| TalAI | + | - | + | - |
| TalAL | + | - | + | - |
| TalAO | + | N* → NG | + | N* → NG |
| TalAP | + | - | + | - |
| TalAQ | + | - | + | - |
| TalAR | + | - | + | Last 3 RVDs missing (truncated) |
| TalAS | + | N* → NG | + | - |
| TalBA | - | | - | |
| TalBJ | + | N* → ND | + | N* → NG |
| TalBK | + | - | + | - |
| TalCA | + | HG → H*, NN → N* , N* → HN | - | |
| TalIT | + | - | + | - |
| TalJR | TalJS | N*→NG, N*→ HN | TalJS | N* → NG, N* → NG |

**Supplementary Table C.** Number of ONT reads completely spanning individual TALE clusters (+ 1 kbp upstream and downstream) for each of the sequenced *Xoo* and *Xoc* strains.

[Available as separate file Frequency_of_reads_in_TALE_regions.xls]

#### Supplementary Files

**Supplementary File S1.** Archive of the progressiveMauve alignment of the different assembly variants presented in Figure 3 for interactive exploration.

[Available as separate file interactiveMauvePXO35_Assemblies.zip]

#### Supplementary Methods

##### Preparation of Beads Solution

Unwashed MNPs were vortexed thoroughly for 5 min. The required amount of beads (e.g. 200 µl for 2 ml of beads solution) was transferred to a 2.0 ml reaction tube, 1 ml of ddH2O was added and the mixture thoroughly vortexed. The tube was then placed on a magnetic rack. After 1 min, the supernatant was removed, and another 1 ml of ddH2O was added. This was repeated 3 times, or until the supernatant becomes clear after all MNPs were stuck to the tube wall when placed on a magnetic rack. To prepare the beads solution (10 mM Tris-HCl, pH 8.0; 1mM EDTA; 1.6 M NaCl; 11 % PEG 8000 (w/v); 0.2 % Tween 20 (v/v); 10 % washed MNP´s (v/v)), all compounds, without the washed MNPs and the PEG were mixed in a 1.5 ml tube. 1 ml of the premixed solution was used to resuspend the washed MNPs and transfer them to the 1.5 ml tube. In the end, the PEG was added, using a cut 1000 µl pipet-tip. All compounds were autoclaved as stock solutions before used.
